# Millimetre wave radiation activates leech nociceptors via TRPV1-like receptor sensitisation

**DOI:** 10.1101/480665

**Authors:** S. Romanenko, A. R. Harvey, L Hool, R. Begley, S. Fan, V. P. Wallace

## Abstract

Due to new applications such as wireless communications, security scanning, and imaging the presence of artificially generated high frequency (30-300 GHz) millimetre-wave (MMW) signals in the environment is increasing. Although safe exposure levels have been set by studies involving direct thermal damage to tissue, there is evidence that MMWs can have an impact on cellular function, including neurons. Earlier in vitro studies have shown that exposure levels well below the recommended safe limit of 1mW/cm^2^ cause changes in the action potential (AP) firing rate, resting potential, and AP pulse shape of sensory neurons in leech preparations, as well as alter neuronal properties in rat cortical brain slices; these effects differ from changes induced by direct heating. In this paper we examine continuous MMW power (up to 80 mW/cm^2^ at 60 GHz) and evaluate the responses in the thermosensitive primary nociceptors of the medicinal leech (genus *Richardsonianus Australis*). The results show that MMW exposure causes an almost two-fold decrease in the threshold for activation of the AP compared with conductive heating (3.6±0.4 mV vs. 6.5±0.4 mV respectively). Our analysis suggests that MMW exposure mediated threshold alterations are not caused by enhancement of voltage gated sodium and potassium conductance. Moreover, it appears that MMW exposure has a modest suppressing effect on membrane excitability. We propose that the reduction in AP threshold can be attributed to sensitization of the TRPV1-like receptor in the leech nociceptor. *In silico* modelling supported the experimental findings. Our results provide evidence that MMW exposure stimulates specific receptor responses that differ from direct conductive heating, fostering the need for additional studies.

## 1. INTRODUCTION

Applications of artificially generated millimeter-wave (MMW) radiation (30-300 GHz) are growing rapidly. Point-to-point wireless communications links, local area networks, security screening systems, and even non-lethal crowd control weapons, have all entered the marketplace. As a consequence human exposure to MMWs is increasing.

Most studies into the effects of MMW exposure on biological tissues conclude that the observed impacts are strictly thermal in nature (1-3). The primary organ which absorbs most of the energy at these frequencies is the skin. Despite that the specific absorption rate (SAR) and power density are maximal at the epidermis, up to 60% of the energy reaches the dermis, and approximately 10% impacts the subcutaneous hypodermis (4). Early studies concluded that MMW radiation has no detectable pathological impact on skin cells and does not cause carcinogenic or other potential long-term effects (1, 5-7). Nevertheless, at least one study concluded that approximately 5% of the observed long-term exposure effects of MMWs is caused by the electromagnetic field interaction with the tissue and not just the thermal impact. (8).

A few studies have looked at the effect of MMW radiation on free nerve endings located in the skin, as well as on sensory neurons, and especially nociceptors (9-11). In one report (9), the thermal pain threshold upon skin heating with intense MMW radiation (94 GHz, 1.8 W/cm^2^) was approximately 44 °C, which is close to the normal physiological value. In contrast, comparatively low intensity MMW exposure (up to only 14 mW/cm^2^) with little ensuing tissue heating, was reported to have an analgesic/hypoalgesic effect (10, 11). These two examples point to the complexity of mechanisms underlying MMW interactions with tissue, and various effects they have at different levels of exposure.

Retzius neurons of the medicinal leech are naturally active with an approximate spiking frequency of 1 Hz); previous studies found clear changes in their spiking activity when they were exposed to very low power MMWs (<1 mW/cm^2^), (12-15). In addition, when MMW irradiation of the neurons was compared to conductive bath heating with a similar temperature rise (<1 °C), the MMWs caused a repression in spiking or action potential (AP) firing rate, whereas conductive heating led to an enhancement of neuronal activity (12-14, 16, 17).

In this paper we extend these earlier studies by examining the direct effect of higher MMW power (>100 mW/cm2) on sensory neurons in the medicinal leech (neurons that are only active when exposed to a noxious stimuli, such as pressure or heat). Because of high water content in biological samples, the higher power MMW radiation inevitably causes sample heating. One of the most sensitive types of neuron is the thermosensitive nociceptor which has a step-like activation when exposed to thermal stimuli. The response of these neurons is due to the presence of transient receptor potential vanilloid (TRPV) channels in the cell membrane. Considering this, the null hypothesis is that MMW mediated heating would cause the same activation of thermosensitive nociceptor as conductive heating. The goal of this work was to determine the effects of MMW radiation on primary nociceptors and whether they can be accurately modelled.

## 2. MATERIALS AND METHODS

### 2.1. Biological model

The experimental model used here was the leech thermosensitive nociceptor (lateral – Nl), these are sensory receptors in somatic structures which convey nociceptive (pain) information to the central nervous system. The leech nociceptive receptors are remarkably similar to vertebrate nociceptors (18), and provide numerous experimental, procedural advantages. Figure 1 shows the relationship of nociceptors to the rest of the nervous system. An advantage of this model is the ability to study individual nociceptors within a whole intact neuronal ganglion, which preserves interactions with other neurons and surrounding glial cells, giving a more representative “whole organ” response.

**Figure 1.**
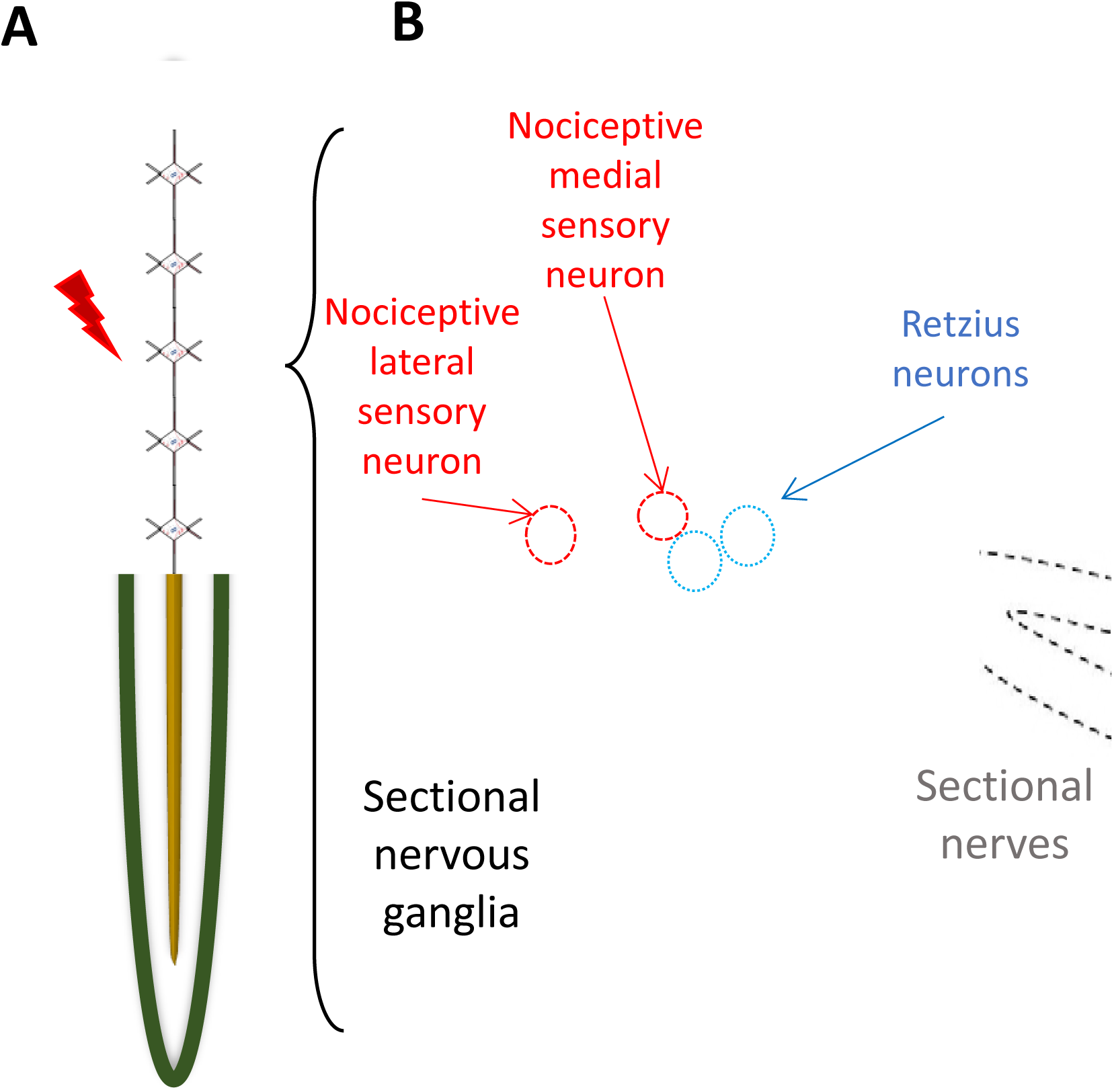
Schematic representation of pain transduction in the leech: (A) Schematic representation of leech body (lower part) and leech segmental nervous system (upper part). Application of supra-threshold thermal stimulus (dark red flash arrow) to the body wall causes activation of the N lateral primary nociceptive neuron. (B) Photograph of single sectional ganglia with lateral nerves (dashed line) and intersectional axonal bundles. Nociceptive neurons depicted in red. Retzius neurons are depicted in blue.

Adult medicinal leeches, genus *Richardsonianus Australis*, were obtained from the Genki Centre, Glebe NSW, AU. Groups of 25 leeches were kept in a glass aquarium with artificial pond water (36 mg/l Instant Ocean salts; Aquarium Systems, Mentor, OH, USA) in a temperature controlled room at 18°C and with a 12 hr light/dark cycle. At the time of the experiments, leeches weighed between 1 and 3 g. Before dissection, the leech was anesthetized in ice cold saline (leech saline) containing: 115 NaCl, 4 KCl, 1.8 CaCl_2_, 1.5 MgCl_2_, 10 HEPES, and 10 glucose (pH = 7.35), all values in mM (reagents from Sigma-Aldrich, St. Louis, MO, USA). Individual ganglia were dissected from midbody segments (M6–M12) and pinned down in a Sylgard184 (Dow Corning Corp., Midland, MI, USA) filled dissection box. Dissected ganglia (Fig. 1 D and 2 C) were transferred to a Petri dish (d: 35 mm) and pinned down (ventral side up) on a paraffin bed (Fig. 2 B) with four tungsten pins. Between experiments the ganglion was maintained at an ambient temperature of 21–25 °C. In experiments with N-(3-Methoxyphenyl)-4-chlorocinnamide (Sigma-Aldrich SB366791) – a selective TRPV1 antagonist – the stock compound (0.1 M in DMSO stored at −20 °C) was diluted in modified (Ca-free) leech saline to make a 50 µM final solution.

**Figure 2:**
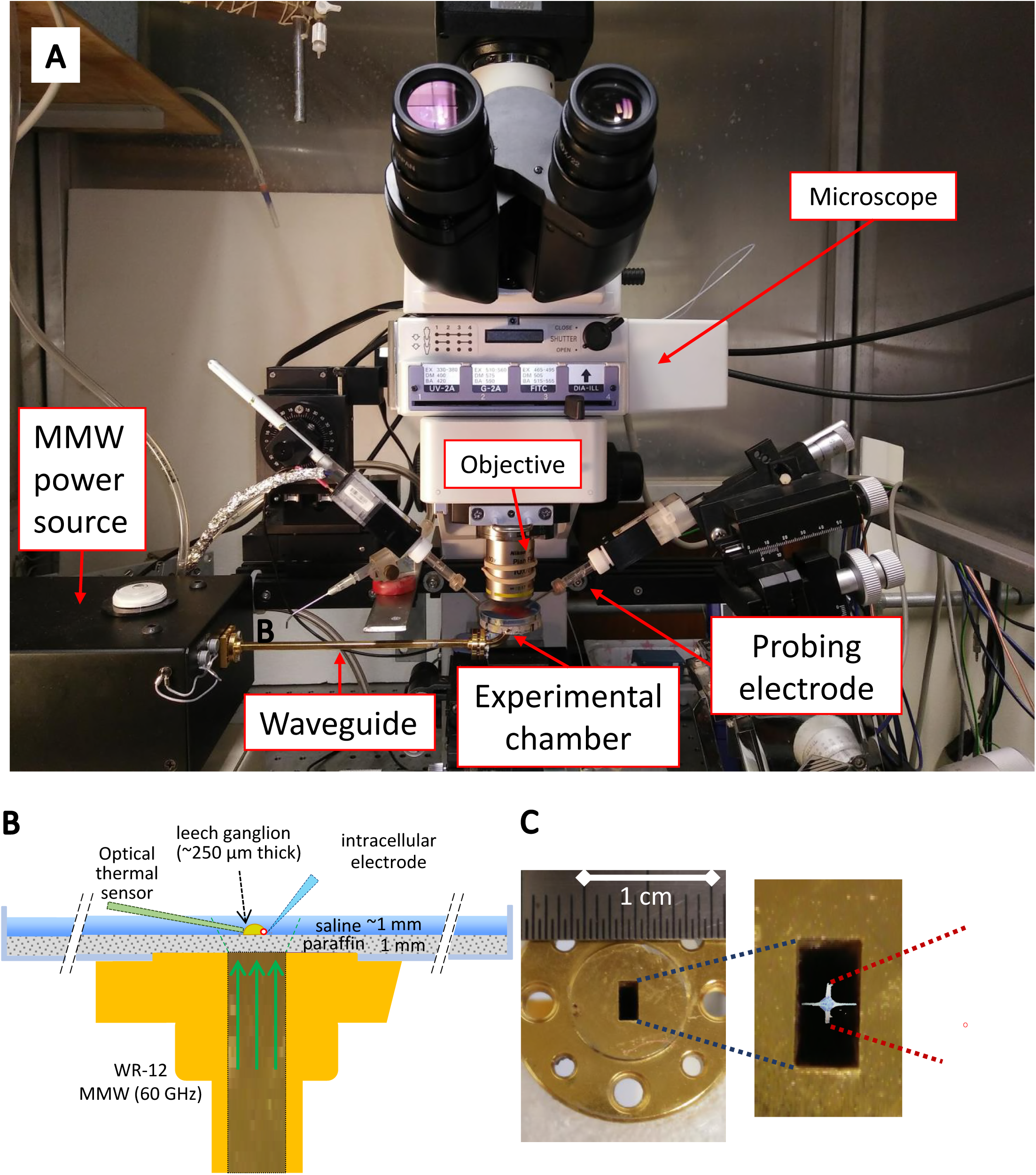
The experimental setup, leech ganglia preparation and experimental chamber is shown. (A) A photograph of experimental setup; (B) Side view of the experimental setup with chamber installed on the WR-12 waveguide, the ganglia is mechanically fixed with fine tungsten pins on the paraffin pad (on the picture the ganglia size is exaggerated for the convenience of the reader). The neuron of interest was probed with a glass microelectrode; a second auxiliary glass electrode was installed as close as possible to the first one and was used for compensation of thermal and convection related artifacts. The reference electrode was mounted on the edge of the chamber to avoid interference with radiation. Thermal sensor was placed as close as possible to the ganglia perpendicularly to the probing electrode; (C) A photograph of WR-12 waveguide top view (leftmost picture) with enlarged picture of its aperture overlapped with a picture of the leech ganglia (middle picture) and enlarged photograph of the ganglia (rightmost picture) stained with 1% Trypan blue, fixed in cold 4% paraformaldehyde 0.1 M phosphate buffer (pH 7.4) and washed in PBS. The red circle indicates the position of one N lateral (Nl) thermosensitive nociceptor neuron.

The Petri dish with the dissected ganglion was placed on an upright microscope (Nikon Eclipse E600-FN, Nikon Corp., Tokyo, Japan) and filled with leech saline so that the ganglion was covered by about a 1 mm-thick layer of fluid. The total volume of saline used was 3 ml. For proper illumination of the ganglion during insertion of the electrolytic probe into a specific neuron, we used a bright white LED at a very oblique incidence angle with respect to the microscope bore (19) and a 10X Plan Fluor objective with differential interference contrast (DIC) filters.

### 2.2. Electrophysiology

To perform the electrophysiological recordings from neurons, a sharp (<1 μm diameter) intracellular electrode was fabricated from aluminosilicate capillary glass (0.87 x 1.2 mm inner/outer diameter, AF120-87-10, Sutter Instruments, CA, USA) using a laser micropipette puller (Sutter Instruments P-2000). The electrodes were filled with 3 M K-acetate and 20 mM KCl unbuffered solution. Average resistance was in the range of 24–27 MΩ. All electrophysiological recordings were performed using a microelectrode amplifier (Axoclamp 900 A; Molecular Devices, Sunnyvale, CA, USA) in the current-clamp mode, held at 0 nA. Recordings were then digitized at 100 kHz using data acquisition hardware (Molecular Devices DigiData 1550 A) and custom software (Molecular Devices Clampex 10). Electrophysiological recordings were performed using the “gap-free” mode and episodic stimulation, and were processed using Molecular Devices Clampfit 10 software. The results were evaluated for two types of sample heating: MMW irradiation and conductive heating of the bath. Details for the MMW irradiation are given in the MMW setup section which follows. Conductive heating of the sample was performed using a 68W 40×40 mm Thermoelectric (Peltier) Module (TEC1-12708 / ZP9104, JayCar Electronics, NSW, AU) with the sample dish installed on top of the module (no saline perfusion was performed). To separate the ganglion from the hot plate, a 2 mm thick paraffin layer was spread across the bottom of the sample. The ganglia were stretched out and pinned directly to the paraffin as shown in Figure 2 B. The heater was positioned off centre from the ganglia so that MMW power could be delivered directly to the cells from below, without having to pass through the Thermoelectric Module (see MMW setup).

The temperature of the saline in the conductive heating experiments was monitored with a type K thermocouple having an accuracy of 0.1°C (QM1283, JayCar Electronics, NSW, AU). Additionally, to avoid any influence of the MMWs might have on the temperature measurement a GaAs crystal based fiber-optic thermometer with a resolution 0.01 °C (OTG-M280 sensor with PicoM spectrophotometer PCM-G1-10-100ST-L; Opsens, Québec, Canada) was used for all of the MMW experiments; the equivalence between the two methods of recording temperature was confirmed (with an accuracy of 0.1°C).

### 2.3. Cell viability experiments

The sample chamber used for the electrophysiological experiments was also used for fluorescence experiments to maintain identical environmental conditions. The Nikon Eclipse microscope was equipped with an epifluorescence attachment (Y-FL), and a high-pressure Mercury lamp (USH-102DH, Ushio Electric Inc., Japan) was used for illumination. Samples were placed under the 10X Plan Fluor DIC objective and after MMW exposure (or sham experiment) the saline in the Petri dish was quickly replaced with a propidium iodide / leech saline solution. The pinned out ganglia were incubated for 60 seconds afterwhich fluorescence images were acquired. The propidium iodide solution was made from a stock solution (Sigma-Aldrich P4170; 1 mg/ml stored at −20°C) by adding 1 µL stock to 1 ml of leech saline. The fluorescent images were collected by a digital CCD camera (DS-5Mc, Nikon, Japan) and stored. Resulting images were processed and analysed with freely available ImageJ software (http://imagej.nih.gov/ij/) (20).

### 2.4. MMW setup

The MMW exposure system consisted of a tunable microwave source from 8-20 GHz (YIG tuned oscillator SAO 002, Virginia Diodes Inc. Charlottesville, VA, USA), coupled to a frequency multiplier/power amplifier chain (Virginia Diodes WR12AMC-HP) which delivered continuous MMWs in the range 60-90 GHz with a maximum output power at any frequency of 100 mW (Fig. 2 A).

The MMW signal was coupled into the ganglion through a single mode open-ended rectangular waveguide (WR-12 Instrumentation Grade Straight Waveguide with UG-387/U Flange Operating from 60 GHz to 90 GHz with a 3.09 × 1.54 mm aperture) placed directly under the Petri dish and encapsulated paraffin layer (Fig. 2 B, C). Using the microscope, the ganglia was aligned directly above the centre of the waveguide aperture, where the MMW field is at a maximum (Fig. 2 C). Note that the MMW power is incident on the ganglia after passing the thin layer of paraffin above, and not through any significant amount of saline, which has a very high MMW absorption coefficient. Both the paraffin and polystyrene have extremely low loss tangents at these wavelengths.

### 2.5. Dosimetry and Radiation exposure simulations

The attenuation of MMW power from the exit port of the waveguide to the ganglion was measured using a pyroelectric detector system equipped with a broad band Winston cone (multimode power collecting funnel) and an infrared blocking filter (DLA-TGS with a 6 THz low-pass edge filter; QMC Instruments Ltd., Cardiff, UK). With no saline present, the attenuation of the MMW power through to the top of the paraffin was 43%, with respect to the power at the waveguide output. When the full 3 ml of saline was added to the Petri dish (to a level 1 mm above the paraffin) the MMW signal was reduced by 31 dB (<1/1000^th^ of the power at the waveguide output) due to the high absorption by water.

To estimate the MMW power density across the ganglion, one must know both the absorption and reflection coefficients of the intervening layers between the ganglion and the waveguide aperture as well as the radiation divergence that occurs as the MMW energy exits the waveguide aperture and is then refracted at each intervening surface. This had to be estimated using an electromagnetic (EM) model of the experimental setup, since the pyroelectric detector was not able to probe the MMW fields at the length scale of the ganglia (generally smaller than one wavelength). A commercial EM simulator based on finite difference time domain (QuickWave; QWED, Warsaw, Poland) (13, 14) was employed for this purpose. Distribution of the MMW power density along the vertical axis, and at selected horizontal cross sections, were estimated from an EM simulation of the exact layout of the waveguide and Petri dish at 60 GHz. The EM simulator requires prior knowledge of the relative permittivity ε_r_ and conductivity σ of all materials encountered. Values for the leech saline solution, the paraffin and polystyrene Petri dish were measured using a commercial THz time-domain spectroscopy (THz-TDS) system (TeraPulse 4000, TeraView, Cambridge, UK) over the temperature range used in the experiments (24-45 °C). At ambient temperature the measurements yielded: saline ε_r_ = 12.01, σ = 72.64; the paraffin padding ε_r_ = 2.27, σ = 0.02; polystyrene ε_r_ = 2.56, σ = 0.09. The properties of saline increase linearly with temperature as expected for an aqueous solution (21). For the leech ganglion, the dielectric constant and conductivity were estimated to be the same as those measured brain tissue samples in the literature (22) of ε_r_ = 10.9 and σ = 48.5 were employed for the EM simulations.

EM simulations were performed at 60 GHz with 100 mW of continuous wave power exiting the waveguide aperture. The calculated power density distributions at various positions along the path from waveguide aperture to ganglion are shown in Fig. 3. The power density varies across the waveguide aperture as shown in the cross section and is 470 mW/cm^2^ at the center of the guide. Beam expansion modified by refraction and reflection at each surface and absorption within each dielectric layer accounts for a difference in power density of approximately 17 % from the waveguide aperture to the top of the ganglion. Accordingly, the power density at the center of the ganglion is 82 mW/cm^2^, while at the bottom is was 170 mW/cm^2^. Note that the power distributed across the diameter of the ganglion is uniform so that any neuron that is probed in the actual experiment receives the same MMW energy and have the same SAR. Also note that the saline surrounding the sides and above the top of the ganglion absorbs all the remaining incident MMW energy so that reflections from the top of the saline layer can be ignored. It should be noted that the IEEE safe exposure limit adopted by the U.S. is 1 mW/cm^2^ for a 6 min exposure (23, 24).

**Figure 3:**
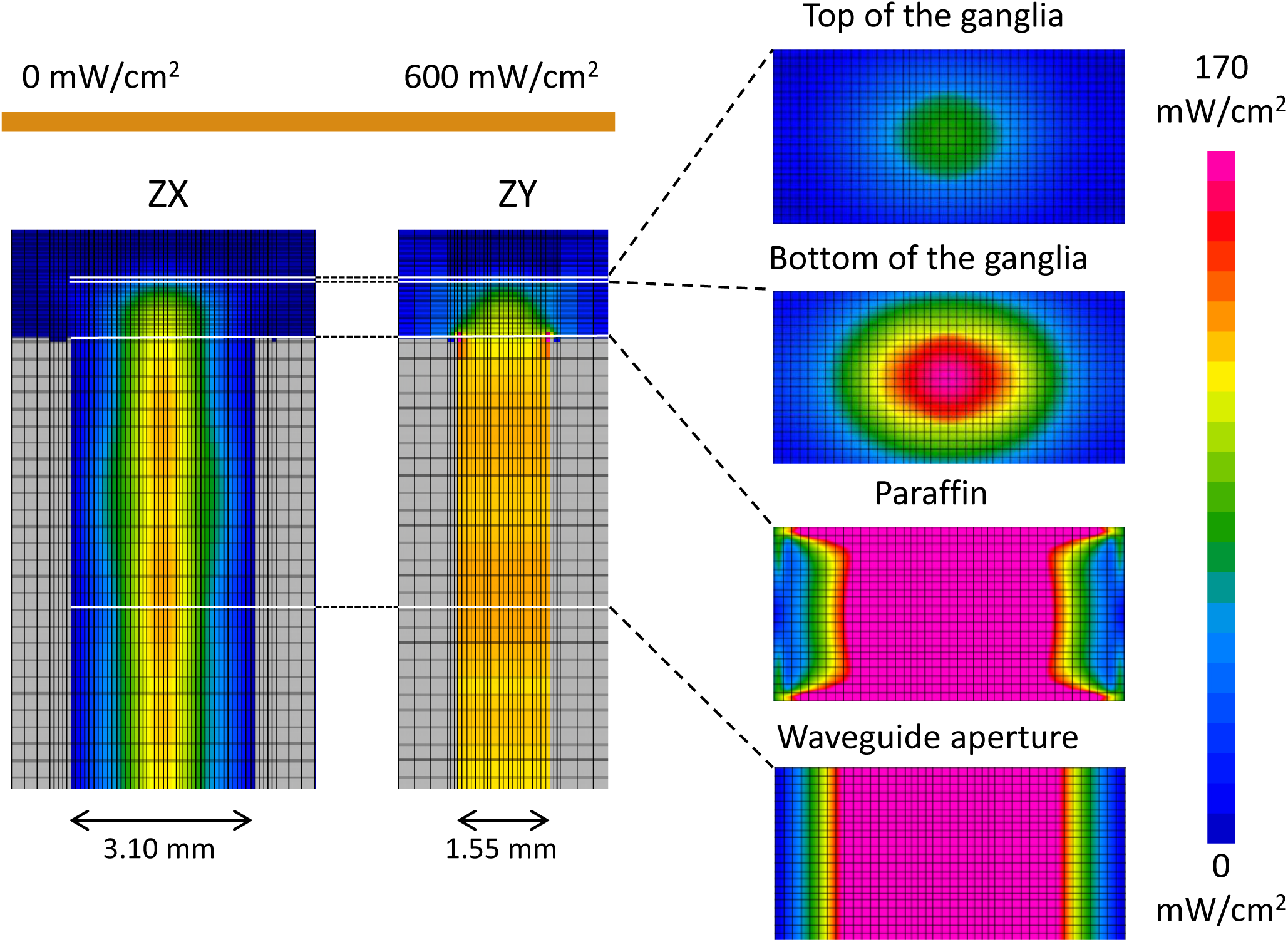
FDTD simulation of MMW power distribution in the experimental setup. Power distribution is colour coded with reference bar provided. Power distribution throughout the waveguide and chamber in ZX and ZY projections (on the left) and power distribution on the different XY levels of the model (on the right). Details in the text.

### 2.6. Biophysical Simulations

Simulations of the electrical activity of the leech lateral N-cell (Nl) were performed with NEURON Simulation Environment (version 7.4, Hines, Yale Univ., USA) (25-27). The neuron model was inspired by the work of Nicholls and Baylor (28) and was modified from Baccus (29) and includes voltage-gated sodium TTX resistant (I_NaTTXR_), and TTX sensitive channels (I_NaTTXs_); potassium delayed rectifier (I_Kk_) and potassium transient current (I_A_) channels; calcium current channels (L-type-like); calcium-dependent potassium current channels (I_KCa_); Na/K –ATPase (I_Na/K-ATP_); leakage current (I_L_); mechanisms for intracellular calcium and sodium accumulation (Ca –accum, Na -accum); and the plasma membrane Ca^2+^ ATPase (PMCA – Ca-pump), and thermosensitive receptor – vanilloid receptor type 1 – TRPV1 channel (I_TRPV1_).

Modeling of the Gouy-Chapman-Stern (GChS) potentials and ion distribution in near proximity, and on the neuronal membrane surface, was performed using formulations provided by (30) and (31). The model was modified to take into account the temperature dependence of the ion hydrated radii (32) and concentration dependent component adapted from Stogryn (33). It also includes the thermal dependence of the leech saline dielectric permittivity measured with the THz time-domain spectroscopy system. Values for the leech saline solution at different temperatures were measured using TeraPulse 4000 in combination with a variable temperature (heated) sample holder (Electrically heated plate, Specac, Kent, UK). At the highest temperatures reached in the experiments we measured for saline: ε_r_ = 17.87, σ = 87.41.

#### Statistics

The MMW exposure and conductive heating effects were analysed using the statistical packages in OriginPro8 (OriginLab Corporation, Northampton, MA, USA) and MatLab 2015 (MathWorks, Natick, MA, USA). The data sets were evaluated by the Kolmogorov-Smirnov normality test. For normally distributed data, a one-way ANOVA analysis was applied followed by one sided post-hoc Holm-Bonferroni and Tukey’s t-tests for inter-group means comparisons. For non-normally distributed data the non-parametric Kruskal–Wallis test (KWANOVA) was used.

Data retrieved from the gap-free electrophysiological recording mode were processed with a modified MatLab code that includes the methods of (34). All codes are available on GitHub. The data were analyzed using a linear regression model (the closeness of the fit is expressed as – R) and a correlation analysis for a series of functions of temperature. Both were performed using standard MatLab functions (solvers *corrcoef, regress*). Significance levels of p < 0.05 were used. The results are reported as mean ± SE (unless otherwise specified).

## 3. RESULTS AND DISCUSSION

### 3.1. Vitality test

All electrophysiological experiments were performed in three mins or less to minimise any potential cell damage. To confirm that the highest intensity MMW exposures did not cause damage to neurons in the ganglia, a viability test was conducted. We used propidium iodide (PI), a DNA binding stain, which does not enter healthy neurons, but passes through the membrane of damaged cells, to assess viability. We used our standard experimental arrangement with saline surrounding the ganglia. The sample (n=11) was rapidly heated from a starting temperature of 22-25 °C to a final temperature of 38-40 °C. After the bath temperature reached its maximum value, the ganglia were kept exposed to MMW radiation for five mins using 100 mW at the waveguide port, and approximately 170 mW/cm^2^ at the bottom of the ganglia. At the end of the exposures, the saline in the measurement chamber was replaced with a PI solution at 22-25 °C. Fluorescent images of ganglia were taken within one minute of the MMW exposure and luminescence was measured with a photometer in the microscope eyepiece. The samples exposed for five mins showed no difference from the control (n = 5 vs. n=7 for MMW vs. Sham exposure). However, the samples exposed for more than 15 minutes (n=6) had a noticeable accumulation of PI in the neurons and the microglial cells located on the surface of the ganglionic sheath. The estimated median lethal dose, LD50 = 8.21 min. We therefore concluded that for heating and exposure times below 5 min, there was no evidence of cell death.

### 3.2 Electrophysiological responses of Nl neurons exposed to MMWs compared to conductive heating of the bath

#### 3.2.1 AP Voltage Threshold

The major thrust of this work was to determine the impact of MMW exposure compared to conductive bath heating on sensory neurons, specifically on the voltage activation threshold. We used a standard Axoclamp 900A and K-acetate-filled pulled capillary microelectrodes to probe changes in the electrical activity of Nl neurons in the medicinal leech. These nociceptive neurons are known to respond to a wide range of stimuli including mechanical, chemical, osmotic and, most important for our study, temperature changes. The NI thermal sensitivity is attributed to the presence of TRPV1-like capsaicin receptors (35). It is well known that TRPV1 are polymodal ionotropic receptors, and in mammalian cells, activate at a thermal threshold in the range of 42-45 °C (36-39). In leeches, the activation threshold is somewhat lower in temperature (40); our experiments confirmed this lower thermal threshold by demonstrating stable and reliable activation of Nl neurons at 38 °C (Fig. 4). In all our experiments, both conductive bath heating and MMW irradiation activated Nl neurons and caused the formation of AP trains. The NI neuron responses to both convention heating and MMW irradiation were recorded in real time, and under identical conditions. Generally, the Nl neurons have a high threshold action potential (AP) and rarely generate spikes if no stimulus is present. Once activated, however, they generate a continuous AP train as long as the stimulus is present (Fig. 4 A, B). Although, the pattern of response is somewhat differed. After being activated the return of the neuron to its initial non-excited state is different between MMW and conductive heating. The AP firing for the MMW exposure cease quickly after the MMW power is turned off, even though the bath temperature is still above the activation threshold. Whereas when the NI neurons are exposed to conductive heating, the AP firing continues even though the bath temperature falls well below the original threshold (show hysteresis-like effects).

**Figure 4:**
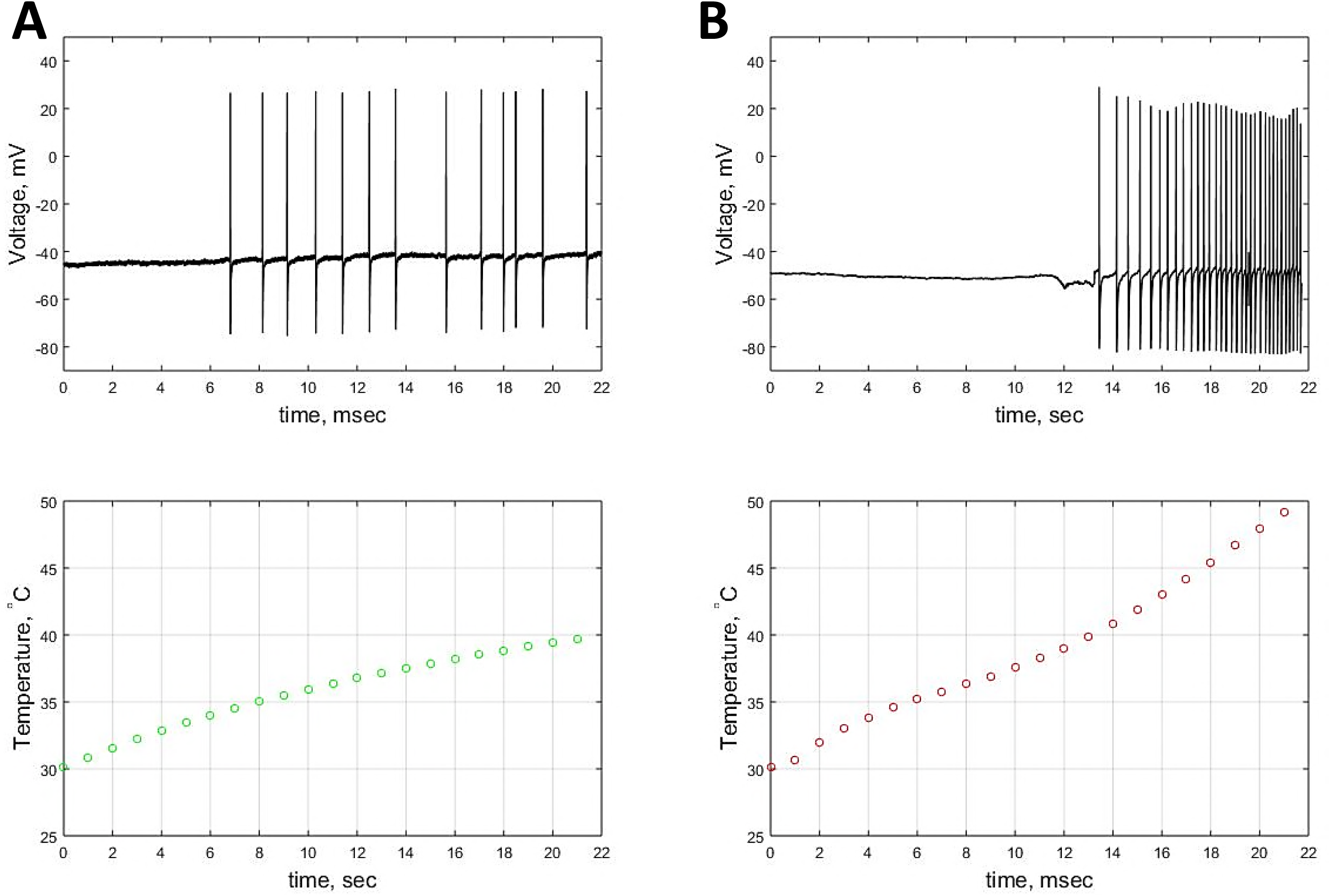
The effect of MMW and conventional conductive heating on Nl neurons: (A) The representative records of Nl neuron activation by thermal stimulus using MMW irradiation. Upper trace is the transmembrane voltage recorded from Nl soma, lower trace is the temperature of surrounding saline. (B) The representative records of Nl neuron activation by conductive heating. Notice, under conductive heating the neuron AP fire frequency increases with the saline temperature increase.

The thermoelectric hotplate was used to generate a temperature increase of equivalent magnitude as that due to the millimeter-wave exposure. In both cases (MMW exposure and conductive heating) the entire neural ganglion is exposed to the same thermal stimulus, since in the case of the MMW exposures, the power density in the exposing beam is uniform across the ganglion.

We defined an activation threshold if at least five AP signals are generated in a train. The results of our experiments on AP activation are summarized in Figure 5. The mean voltage threshold of AP activation for conductive heating was 6.50±0.36 mV (n = 21), whereas for MMW exposure it was 3.63±0.38 mV (n = 17; p<0.001). The voltage threshold for AP (throughout the entire range of observed temperatures) from conductive heating activation varied from 0.5 to 19.5 mV, with two distinct, and very substantial, peaks at 2.78±2.26 mV and 9.84±7.20 mV (Fig. 5 B) – obtained from fitting with Gaussian function, whereas data obtained from the MMW irradiated samples shows no such strong variation with temperature (1.83±1.36 mV). We do not have a good physiological model for this behavior, however others have noted similar enhanced AP firing due to activation and inactivation kinetics of ion channels, conduction velocity, and synaptic events amongst other phenomena (41-43).

**Figure 5:**
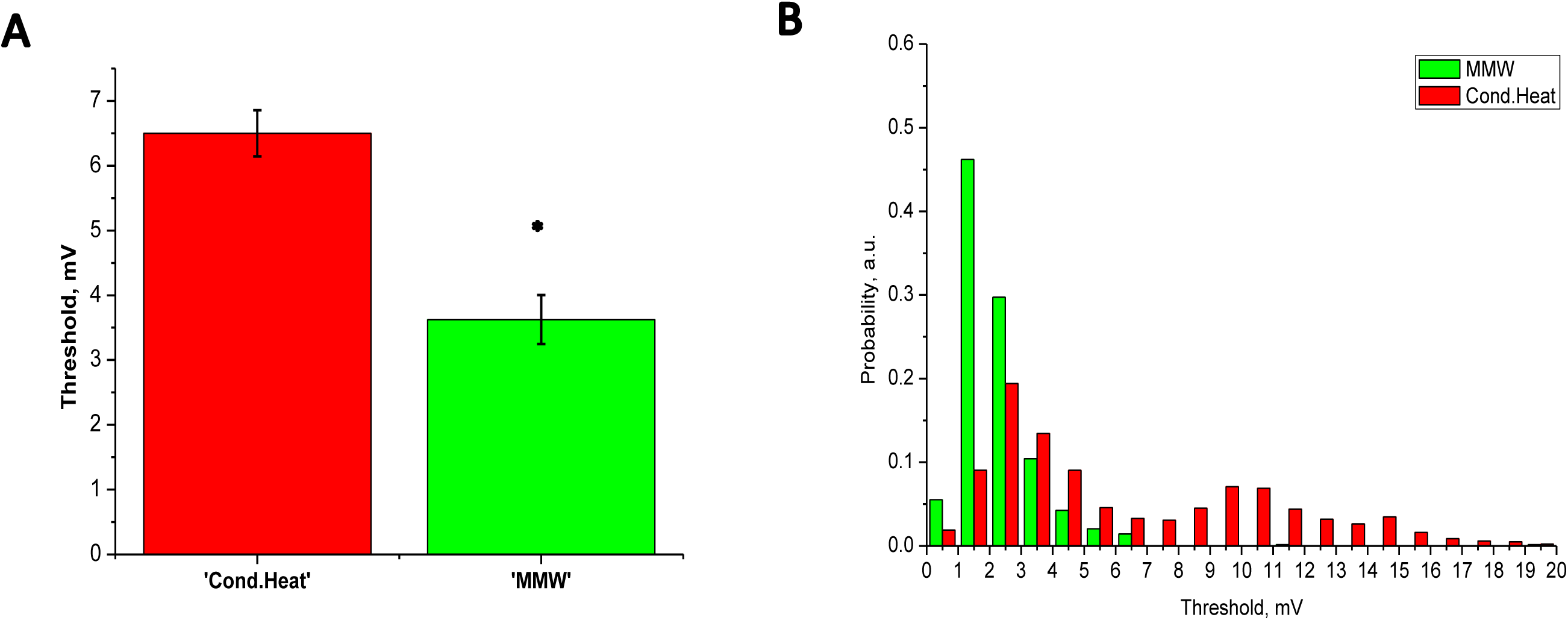
Effect of conductive and MMW heating on Nl neuron activation threshold: (A) The Nl neuron voltage thresholds averaged over first 5 AP in a train under the condition of MMW exposure and conductive heating (Cond.Heat). Data are mean±SE; (B) Voltage threshold probability distribution for MMW (green) and conductive (red) heating. Note bi-component distribution for conductive heating.

Interestingly, the threshold temperature of Nl neuron activation was lower for MMW radiation,∼ 36-38 °C (Fig. 4 A vs B) compared with convetional sample heating ∼ 40-42 °C. However, due to the size and location of the temperature probe we are not certain that the temperature of the neuron within the focal region of the MMWs was at a higher temperature, closer to that of the conventional heating.

We studied the correlation coefficients between an assortment of AP parameters and changes of sample temperature for MMW irradiation and conductive heating. The strongest correlation in the conductive heating experiments is found for changes in AP amplitude (−0.80), while the same parameter under MMW irradiation has a lower correlation coefficient −0.68. In general, there is only modest correlative behaviour for the parameters studied (firing rate, AP half-width, rise and fall times) with linear rise in temperature.

Comparison of MMW irradiation and conductive heating as the trigger indicate that neurons subjected to the MMW have a lower activation voltage threshold for AP formation.

#### 3.2.2 AP Current Threshold

In addition to the experiments on AP threshold and observed changes in AP magnitude and shape characteristics, we looked at neuron current stimulus excitability under MMW and conductive heating. We employed a current-clamp step protocol to build up strength-duration curves (Fig. 6 A, B) for a control (22-24 °C) and conditions under which the temperature of the neurons was raised to 31-33 °C by MMW irradiation and conductive heating. The thermal limit was set to keep neurons below the threshold temperature for AP thermal excitation (to prevent any stray AP). A current stimulus of varying amplitude (0-1.4 nA) and fixed duration (55 msec) was injected into the neuron while we recorded the time to the start of AP formation (taken to be the time of the first AP peak). Fig. 6 A and B show the plots of stimulus current versus elapsed time to start of the AP (strength-duration) for a number of measurements (n = 13 vs. 7 for MMW vs. conductive heating respectively). We expect thermosensitive neurons to have nearly zero average rheobase at subthreshold temperatures, and that any added inward current would trigger AP formation. However, the neurons’ response was somewhat different for different types of heating. The entire set of data were fit to (At+B)/t where A is the rheobase – the minimum current at which an AP is triggered if impulse is infinite – and B minimizes the deviation from the fitted line. The estimated average of rheobases obtained from individual fitting of every experimental strength-duration curve were 0.10±0.03 nA, 0.01±0.27 nA and 0.06±0.04 nA for control, conductive and MMW irradiation respectively. As can be seen in Fig. 6 B, there is little difference between the control and the MMW curves. In contrast, the strength-duration curve obtained from experiments with conductive heating is lower than the control curve, as the average rheobase value was near zero.

**Figure 6:**
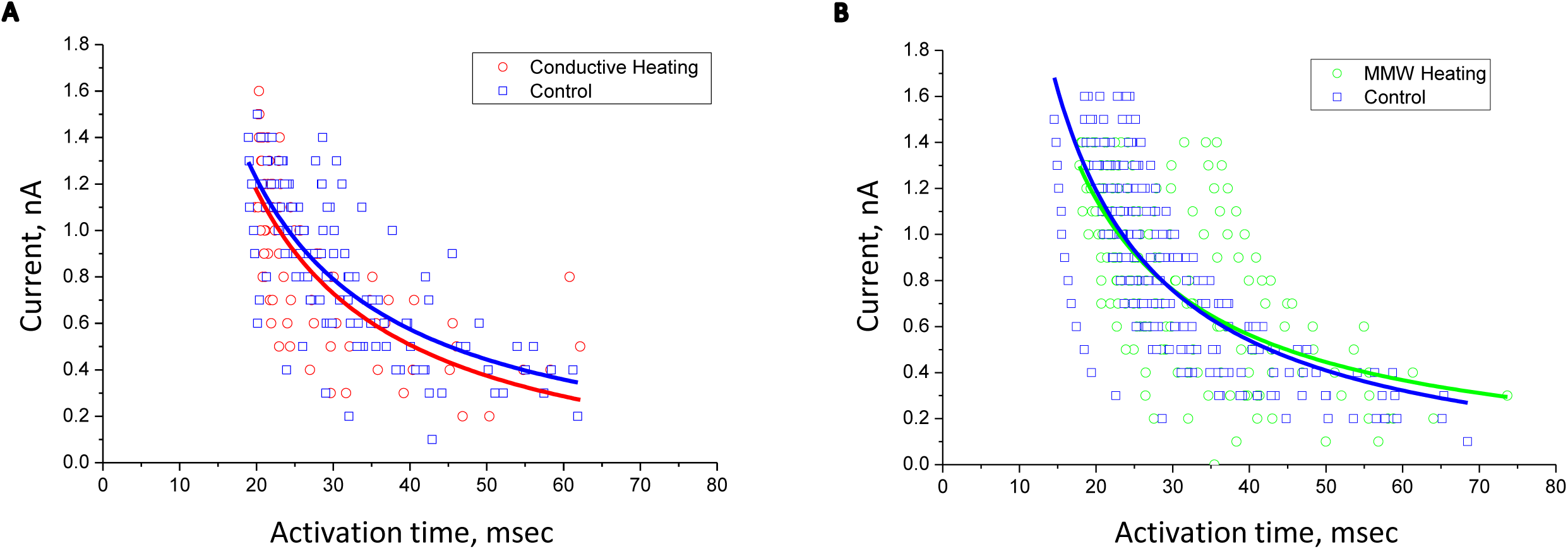
Strength duration curves for NI neuron obtained under control (blue), MMW (green) and conductive (red) heating conditions. Raw data showed in two scatter plots to avoid confusion due to substantial overlapping. Solid line curves are fittings from original raw data with rational function (see the text). Note, in case of conductive heating the curve is below the control one, as would be expected for higher temperatures, whereas for MMW heating there is little if any difference.

#### 3.2.3 Passive parameters

To estimate the contribution from passive electrophysiological properties of the probed neuron we injected negative current pulses that resulted in no AP generation. We then measured the capacitance and conductance under MMW and conductive heating (34 −39 °C) vs the control (20 - 22 °C). The mean electrical resistance was significantly higher in the case of MMW irradiation 68.33±4.10 mΩ (n = 9; ANOVA: p<0.01), than both the control and conductive heating which were 47.54±1.66 mΩ and 52.10±1.90 mΩ, respectively. The capacitance, however had a somewhat lower value for the MMW exposed samples (0.0097±0.004 nF) than either the control or the conductive heated neurons (0.014±0.001 and 0.012±0.002 nF, respectively), although the differences in the three values is considered not significant taking the the errors into consideration. This change in capacitance could be accounted for by a slight shrinkage of the cell size (approx. 10% estimated from cell body). These results show that passive physiological parameters do not change significantly with heating below threshold temperatures, and therefore do not contribute to the differences in the MMW versus conductive heating results we have observed.

### 3.3. Multimodality of MMW effect

#### 3.3.1 The effect of MMW irradiation in the presence of TRPV1 antagonist

As noted in the current-step experiments (Section 2.2), thermal activation by MMW irradiation is not based on changes to the AP driving conductance. However, further work was needed to clarify whether or not changes to thermally sensitive transient receptor potential vanilloid receptor-1 (TRPV1) are the basis for the observed lower voltage thresholds upon MMW exposure. We used a selective cinnamide TRPV1 antagonist (SB366791) to see if this would reduce or eliminate the threshold effects seen under MMW irradiation (44). It is known that this antagonist does not interfere with voltage-gated calcium channels and with hyperpolarization-activated channels (unlike Capsazepine) and it blocks thermal and capsaicin-evoked responses in leech Nl neurons (40). For these experiments, the ganglia were incubated in 50 μM of antagonist for 5 min before being exposed to the MMW irradiation (170 mW/cm^2^ for less than 3 min) and the heating of the bath was limited to 44-45 °C.

Taking the full suite of temperature dependent data and comparing the change in the AP activation threshold (estimated from first 5 AP) yields a significant difference between the neurons in the antagonist and normal saline solutions: 14.75±1.51 mV vs. 3.63±0.37 mV (n = 10, p<0.001) (Fig. 7 B). Also, the neurons in the antagonist solution have an AP voltage threshold probability distribution (Fig. 7 A) that is very similar to that obtained from the conductive heating experiments (section 2.1.) and show a bi-modal distribution as in Figure 6 B. Fitting of the AP threshold distribution using a Gaussian function reveals that there is a good match between the antagonist treated and normal conventional heat activated neurons at both component of bi-modal distribution: 3.75±1.63 mV vs. 2.78±2.26 mV low-threshold component and 19.21±12.94 mV vs. 9.84±7.20 mV for high-threshold component (c.f. section 2.1.). Hence, the effect of MMW irradiation on Nl neurons could be the result of either multi-modal facilitated activation of TRPV1 (simultaneous thermal and osmotic) or activation of another sensory channel in the neuron’s membrane (the option investigated in Section 3.3).

**Figure 7:**
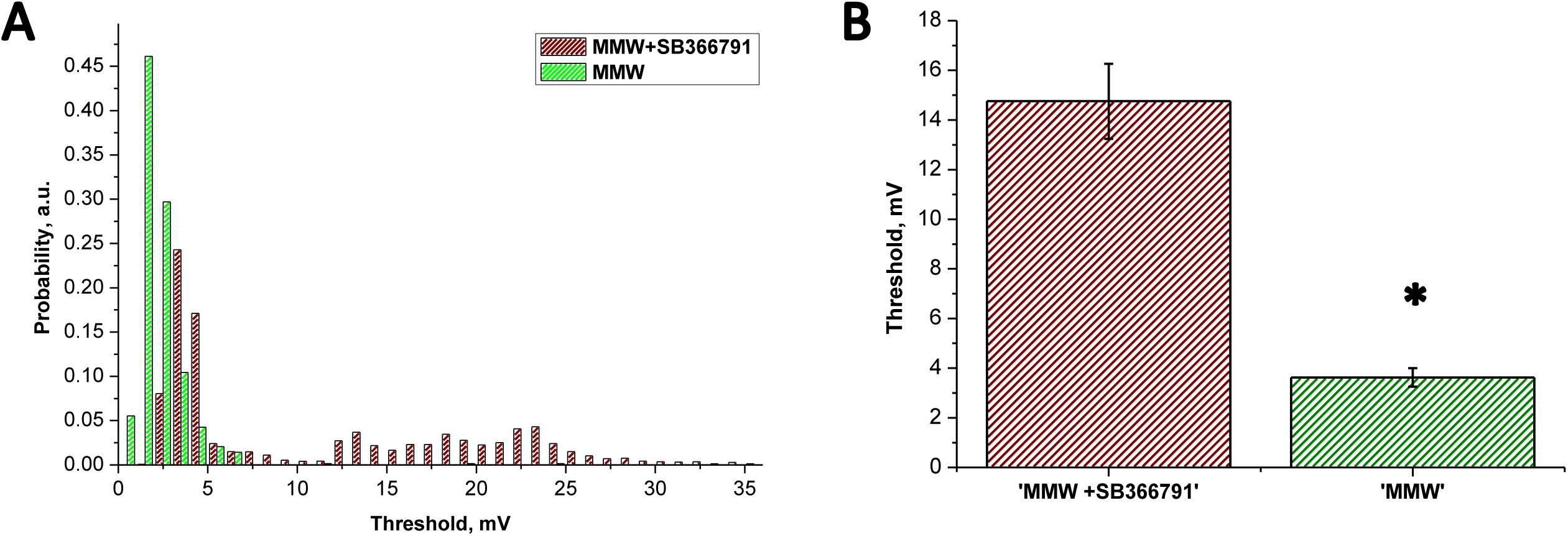
The effect of MMW heating on Nl neuron treated with the TRPV1 selective antagonist: (A) Voltage threshold probability distribution for treated (red) and not treated (green) with antagonist NI neurons activated with MMW heating; (B) The voltage thresholds for Nl neurons treated and not treated with antagonist, (averaged over first 5 AP in the evoked train). Data are mean±SE, p<0.01.

#### 3.3.2 The effect of MMW in the N-medial neurons

To rule out a separate sensory channel, we tested a different type of neuron the N-medial (Nm) neurons in the ganglia which are mechanical nociceptors (45). These neurons are also high-threshold nociceptors and homologs of the Nl neurons. However, they do not express sensitivity to high-temperature stimulus nor do they respond to capsaicin. The Nm neurons were probed in the same manner as the Nl neurons, but in contrast, the introduction of the electrode (causing the mechanical force irritation) results in continuous AP spiking (which is expected for mechanical nociceptors). Even injection of a prolonged hyperpolarizing current did not suppress this activity in the Nm cells. Unlike the NI neurons, MMW irradiation (170 mW/cm^2^ for less than 3 min over a temperature range of 20 - 41 °C) did not result in step-like alterations in neuron activity, but gradual changes were observed. A comparison of voltage thresholds before, and during MMW exposure showed an increase: 17.75±2.36 vs. 23.86±1.79 mV (n=6,; p<0.05) (Fig. 8 A, B). Note, here we compare the AP thresholds from slightly different temperature ranges 21-23 °C for control data and 21-29 °C for MMW data. Even though the sample was heated by MMW exposure the AP threshold under this condition was higher than under control ones. This is consistent with results obtained in Sections 2.2 and 2.3. Since the Nm neuron does not have the thermal sensor (TRPV1 like channel) and hence do not replicate the effect seen for Nl neuron (in section 2.1), we conclude that the mechanism underlying the lower activation threshold in Nl neurons exposed to MMW radiation is due to the multi–modal activation of TRPV1 receptors.

**Figure 8:**
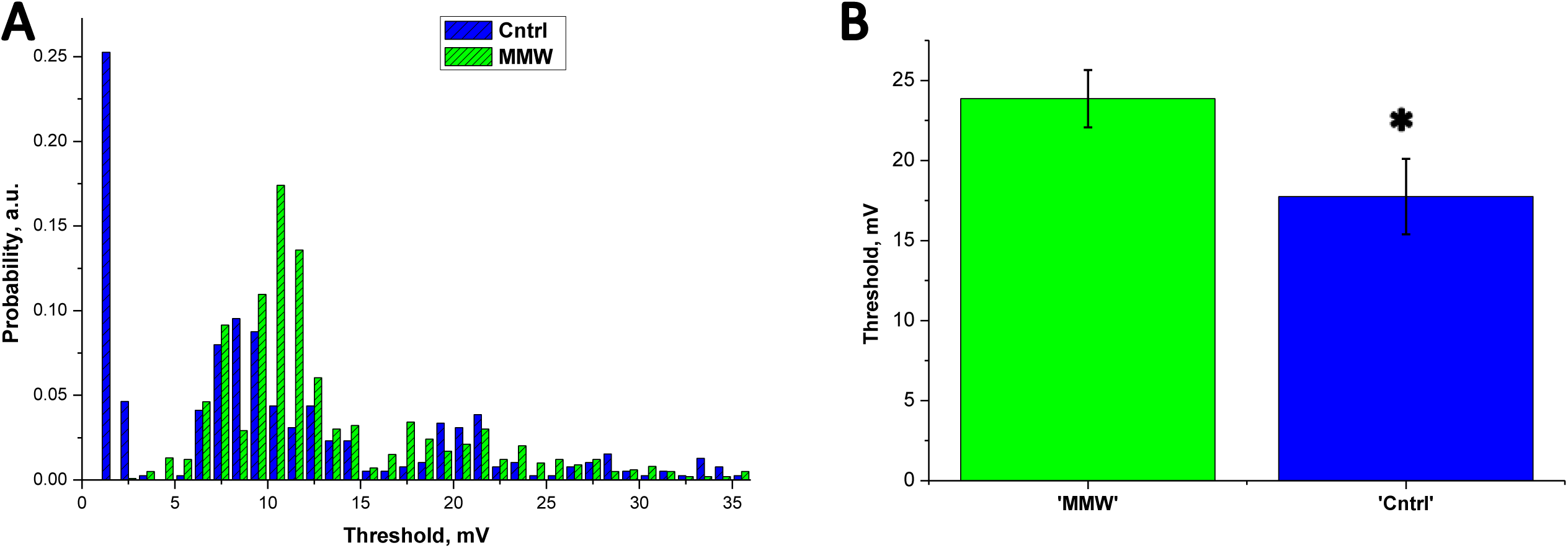
The effect of MMW heating on Nm neuron: (A) Voltage threshold probability distribution for Nm neurons before (Cntrl - blue) and during MMW heating (MMW - green); (B) The voltage thresholds for Nm neurons before (Cntrl) and during MMW heating (MMW), averaged over first 5 AP after MMW application. Data are mean±SE, p<0.01.

### 3.4. Computer simulation of the Nl thermal activation

At this point, we have seen that MMW irradiation of Nl neurons results in electrophysiological changes that are different from exposure to conventional conductive heating. Some differences are pronounced, such as changes to the voltage threshold for AP activation, others were less significant, such as alterations in cell membrane capacitance. To understand whether MMW mediated shifts in Nl sensitivity are a result of simple temperature dynamics or some more complex mechanism, we tested different assumptions on a computer model of the Nl neuron.

Our model was implemented in NEURON simulation environment (version 7.4, Hines, Yale Univ., USA) and is based on neuron simulations such as those reported in (28) and was modified from (29). The model is thermally robust and reliably generates AP’s at simulated temperatures up to 50 °C. The properties of the TRPV1 mechanism were adjusted to replicate the features of leech neurons. The simulation of model Nl neuron activation with thermal stimulus (conductive heating) is presented in Fig. 9 A. After reaching the thermal threshold, the model neuron generates a stable AP train (as observed in experiments). We tested whether the slope and response curve of the temperature rise has any impact on the simulated Nl activation threshold. A comparison of a different rate of heating, similar to those we have measured in our experiments (between 0.7 and 2.3 °C/sec), show no dramatic impact on threshold in the model neuron. We also tested changes in cell capacitance. We took into account that dynamic change of the neuron’s capacitance should be reflected not just in the membrane parameters, but as an additional current (*(dC*_*m*_*/dt)×Um*, where *C*_*m*_ - membrane capacitance, *U*_*m*_ – transmembrane potential) in the AP excitation equation. The rate of capacitance change in the model is taken to be proportional to temperature with a maximum decrease of 10 % compared to the starting value (as it was estimated from neuron fluorescent pictures SI5). The simulations showed no significant change to Nl neuron activation with changing capacitance.

**Figure 9:**
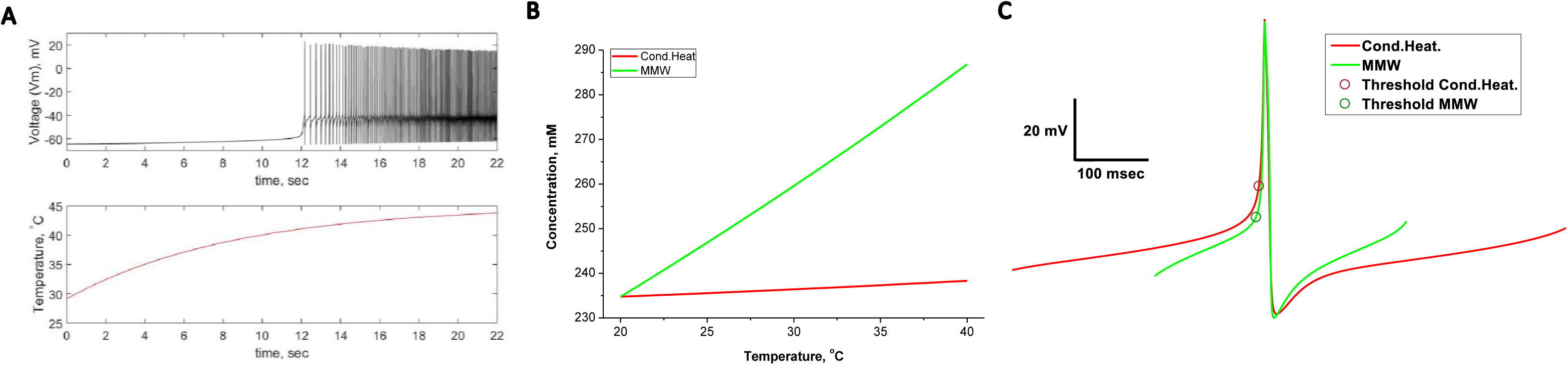
Computer simulation of Nl neuron model (NEURON) and modeling of an ionic double layer on the surface of the membrane (Gouy-Chapman-Stern Model). (A) Simulated traces of model Nl neuron response to a thermal stimulus: transmembrane voltage recorded from cell soma (red), sample’s temperature trace (purple); (B) Calculated with Gouy-Chapman-Stern (G-Ch-S) model changes in monovalent ion concentration on outer surface of membrane against temperature (red) and with concurrent increase (up to 30%) of the surface charge density (green). The red trace reflects the effect of conductive heating, whereas the green trace reflects the effect of MMW; (C) Simulated AP recorded from soma of Nl neuron model for simple temperature increase (red – Cond.Heat.) and with concurrent increase of the surface charge density (green - MMW). Calculated AP activation thresholds are shown on each trace. Note, MMW trace has lower activation threshold if compare with Cond.Heat. one, as it was observed in the real experiment.

From our experiments, we hypothesize that TRPV1 channels are sensitized during MMW irradiation and that two mechanisms, thermal and osmotic activation of TRPV1, are involved. We know from the literature (46) that an increase in osmolality up to 350 mOsm causes sensitization, and can even activate the TRPV1 channel. We assumed that MMW radiation promotes a local peri-membrane increase of solute concentration; however, the question of whether MMWs can change the peri-membrane solute concentration up to 350 mOsm still remains. To address this issue, we used a Gouy-Chapman-Stern (GChS) Model of an ionic double layer on the surface of the membrane (30) (31). The model relates the membrane surface charge with the surface potential and electrolyte concentrations. It accounts for the temperature of the system, ion hydration radii, and the temperature dependence of the saline dielectric properties. Values for surface charge density were taken from the Hille estimation (47-49) – one electron charge per 100 - 400 Å. The asymmetry in charge density for inner and outer leaflets of the membrane were introduced with an outer layer having approximately three times more charge (50). The model calculates the distribution of electrostatic potential and electrolyte concentration within membrane surface proximity. We ran the simulation for different temperatures within our experimental range. Even with a change in temperature of 20 °C the surface potential altered by just a few millivolts. Even smaller changes occur for the peri-membrane concentration of the sodium ions. The relationship between the system temperature and peri-membrane sodium concentration is shown in Fig. 9 B (Cond. Heat). Within the temperature range we tested, the electrolyte concentration shifted only a few mM. Another factor defining potential, and local concentration of electrolyte on the membrane’s surface is surface charge density (*σ*_*out*_, *σ*_*in*_). In contrast to the temperature, a 30% increase in *σ*_*out*_ caused nearly a 10 mV shift in surface potential and a 52 mM increase in surface electrolyte concentration (Fig. 9 B - MMW). Such strong changes could create conditions in close proximity to the membrane that are capable of sensitizing the TRPV1 via the osmosensing mechanism (46, 51). Hence, the combined effect of increased temperature and surface charge density can have an additive impact that alters the TRPV1 activation threshold in Nl neurons. We have calculated the relationship between the local electrolyte concentration and local temperature with concurrent relative alteration in surface charge density (Fig. 9 B - MMW). We find that only a 22 % increase in surface charge density would satisfy the requirements for activating the Nl neuron. These conditions were inserted into our Nl neuron model by increasing the concentration of NaCl on the outer side of the membrane. The result (Fig. 9 C) demonstrates a shift in the AP threshold of Nl neuron activation in same fashion as it was observed in the experiment (section 2.1). The simulation also demonstrated the increase in depolarising inward TRPV1 current caused by MMW type of heating. Hence, among all potential mechanisms underlying the MMW irradiation effect, the most efficient one is simultaneous TRPV1 sensitization via its thermo- and osmo-sensitivity.

## 4. CONCLUSION

This study investigates the effect of MMW radiation on primary nociceptive neurons. By comparing the results of thermal bath heating and via more complex thermal and electromagnetic stimuli present in MMW irradiation, we have tried to elucidate and then assign specific electrophysiological changes to NI activation pathways. Understanding whether this form of radiation (generally not occurring in nature), and its potential effects on primary nociceptors is of vital importance as we move towards a world with dramatically increased presence of these frequencies in the environment.

In our experiments we exploit the natural property of the nociceptors to detect and react to critical levels of thermal stimulus with a strong electrophysiological response. Indeed, thermo-sensitive nociceptors have specific vanilloid receptors (the most prominent being TRPV1) that are capable of detecting thermal stimuli above 38 °C, which we have used in our experimental model. Thus, if the application of MMWs gives rise to a thermal response that deviates from the response caused by conventional heating, it would mean there are MMW specific effects for at least some types of neurons.

Our core investigation compares the stimulation of Nl neurons with conventional bath heating and with radiative heating using MMWs with a power density of 170 mW/cm^2^. We found that irradiating with MMWs facilitates the activation of Nl neurons, with the average voltage threshold for AP formation almost half that of conductive heating alone. We also measured a difference in the temperature of Nl activation (Fig 4) but we cannot define the temperature within the neuron itself as the probe position only measures the surface temperature. Nevertheless, it would be reasonable to expect a decrease in the thermal threshold from a theoretical point of view. Firstly, lowered voltage threshold of AP activation would require smaller activation current carried by thermal sensor within the cell-TRPV1, thus shifting the thermal threshold to a lower temperature. Secondly, MMW mediated sensitisation to TRPV1 itself could lower thermal activation threshold. These mechanisms are to be investigated in future studies.

We ran a series of tests to determine what could be behind this difference. As for many types of sensory neurons (both vertebrate and annelids), the important regulatory mechanism for AP train formation is the calcium-dependent potassium current (K_Ca_). Studies conducted on *Vero* kidney cells exposed to 42.25 GHz MMW radiation (16mW) demonstrated a decrease in open-state probability for K_Ca_ channels (52). Hence, the slow process of depolarisation accumulation after every AP would result in a decrease of the voltage threshold value. However, because of the temperature increase any potential MMW effect on K_Ca_ channels open-state probability would be masked (temperature increase facilitates the movement of voltage sensor).

Heating itself commonly causes a significant increase in AP firing, activation and inactivation of ion channels, changes in conduction velocity and synaptic events, etc. (53-55). In contrast to conductive heating, MMW irradiation does not have such an effect on AP formation, when compared with control data (Section 2.2.). It implies that voltage-gated channels participating in AP formation may not be involved in changes in threshold, but governed by another mechanism; we postulated that this could be the TRPV1-like channel. To test our assumption, we performed an excitability study at temperatures below the heat activation threshold, where TRPV1 is not involved. A comparison of the strength-duration curves for conductive heating and MMW irradiation revealed that conductive heating lead to neuron behaviour as would be expected under varying injected current stimulus – averaged rheobase lowered to nearly zero level (0.01±0.27 nA), whereas MMW irradiation produces no alterations in rheobase when compared to the control (0.06±0.04 and nA0.10±0.03 nA). These results are in agreement with our previous studies conducted on Retzius neurons which have no thermal receptors, and demonstrated some mild suppression of the neuronal activity upon application of MMW irradiation of low intensity (13).

We repeated the MMW irradiation and AP activation with a specific TRPV1 receptor antagonist (SB366791) in the bath solution; to our surprise, Nl neurons still responded to some extent to MMW irradiation. This implies that the activation of Nl neurons is not simply a result of activating the heat sensor of the TRPV1 receptor, but a more complex phenomenon. Indeed, the SB366791 is an effective suppressor of the thermal and capsaicin-evoked responses in TRPV1 expressing cells, but it does not block the sensitivity to some other stimulus modalities (low pH, osmolarity) (44). Moreover, TRPV1 channels possess osmosensitivity properties, and this modality could be facilitated by, or be the facilitator of, thermal and chemical sensitivities (45).

We also tried MMW irradiation of Nm neurons, which did not show any analogue to the neuronal activity alterations observed in the Nl neurons. This confirms the critical role of the TRPV1 receptors in the observed Nl sensitivity shifts. One possibility is that MMWs activate the osmotic sensitivity of TRPV1, and activates the receptor. Indeed, the mean voltage threshold and AP formation probability distributions for conductive heating and for MMW irradiation in the presence of the antagonist are very close in value, and have bi-modal distributions as a function of temperature. In both these cases, only one TRPV1 sensitization pathway is involved in the activation of the receptor – the osmotic one. Whereas, in the earlier MMW irradiation experiments (without SB366791), both the thermal and osmotic sensitization pathways are present. Although MMW exposure does not alter solute concentration in bulk saline, in the peri-membrane space such alterations could occur and our observed effects are in accord with this condition. Correspondingly, the Na/K pump plays an important part in cell volume regulation, and studies with intense 60GHz MMW radiation applied to artificial neurons showed significant up regulation of Na/K-ATPase (56).

To help us better understand the individual contributions from all the observed effects of MMW irradiated Nl neurons, a mathematical model of our Nl neuron was constructed with NEURON simulation software. The model shows the collective effect of all postulated MMW stimulated response mechanisms and separates them from the conductive heating effects on Nl neurons. The model demonstrates that, without the TRPV1 channel, the neuron could not generate AP trains, regardless of the background temperature. Incorporation of a TRPV1 channel with thermal sensitivity only, resulted in the generation of an AP train as soon as the NI temperature crossed the thermal threshold.

Our model shows that the osmotic sensitization of TRPV1 requires only a 17% increase in electrolyte concentration around the cell to cause a significant reduction (79 %) in the Nl threshold. How can MMWs induce this change and why is it not also caused by conductive heating? One possible explanation is that the MMW radiation might stimulate an increase in calcium ion activity, due to a very high specific absorption rate, and as a result, dissociate the calcium from the membrane. Increased *σ*_*out*_ would quickly attract monovalent counter ions, which in the extracellular medium, is mainly sodium cations (Na^+^). The growing concentration of Na^+^ in a peri-membrane space could be sensed by the TRPV1 osmosensor and promote sensitization. Monovalent cations do not compensate surface charge in the same manner as divalent ones, because the binding constants of monovalent cations are 1-2 orders of magnitude less than those of divalent cations (41, 42). It is estimated in our model, that only a 22 % increase in surface charge density would be enough to build up a 17% increase in cation concentration, and as a result activate the TRPV1 receptor. Taking into account that under normal conditions most of the membrane surface’s negatively charged sites are occupied by calcium ions (43), a 22 % increase in surface charge density is plausible. Moreover, an increase in the surface charge density could be partially aided by cell shrinkage, so the dissociation of Ca^2+^ from negatively charged sites could even be smaller. In support of this hypothesis, it has been shown that 42 GHz radiation at a power level of 35 mW/cm^2^ caused translocation of negatively charged phosphatidylserine from the inner to the outer side of the cell membrane (4, 57, 58).

Finally, an increase in *σ*_*out*_ and an increase in the concentration of Na^+^ in proximity to the membrane surface, are in agreement with our data for the step-current protocol for MMW irradiation which are not different to control, in contrast to the observed conductive heating effect. Indeed, studies by Hille (49, 59, 60) demonstrate that increased monovalent ion concentrations promote a positive shift in cell surface potential and thus make membranes less excitable. Other researchers (61) also showed that an increased negative charge on the membrane surface promotes resting state stability.

In summary, this work has demonstrated significant changes in the functional properties of thermosensitive primary sensory neurons subjected to MMW radiation that differ from the impact of direct conductive heating. Nociceptors subject to intense MMW stimuli showed a notable decrease in the voltage threshold of AP formation, compared to conventional conductive heating. Experiments and simulations confirmed the central role of the TRPV1-like channel in the observed phenomena. We conclude that the distinctive impacts of MMW radiation on NI neurons is based on sensitisation of a TRPV1-like channel and that the observed lower thermal nociceptor activation threshold is a result of the combined action of temperature and peri-membrane osmolarity. The latter could be considered as the distinctive non-thermal component of MMW irradiation effects on cells, as this osmolarity sensitization is not present with conductive heating. This two-modality sensitization mechanism is likely transferrable to other types of neurons exposed to MMW radiation. This study may be of use when considering potential safety guidelines for future MMW sources resulting in unregulated public exposure. Furthermore, because MMW radiation can be directed and focused, its effects on AP thresholds may have potential uses in neurostimulation or chronic pain suppression.

## ACKNOWLEDGEMENTS

We are thankful to Dr. Victor Pikov (Bioelectronics unit, GlaxoSmithKline Bioelectronic Medicine) for technical support and the sharing of some equipment with us during the course of this research. We also thank Dr. Peter Siegel (California Institute of Technology) for discussions and reviewing of the manuscript.

## AUTHOR CONTRIBUTIONS

V.P.W. and S.R. conception and design of research; S.R. and S.F. performed experiments; S.R and S. F. analysed the data; S.R. performed NEURON and Matlab simulations; R.B. performed RF FDTD simulation; S.R., V.P.W., A.H., L.H. interpreted results of experiments; S.R. R.B. S.F. V.P.W. prepared Figures; S.R. and V.P.W. drafted manuscript; A.H., L.H., H.M., D.R. and P.S. edited and revised manuscript; S.R., A.H., L.H., P.S., H.M., D.R., R.B., S.F., and V.P.W. approved final version of manuscript.

